# SMCHD1 loss triggers DUX4 expression by disrupting splicing in FSHD2

**DOI:** 10.1101/2023.02.27.530258

**Authors:** Eden Engal, Aveksha Sharma, Nadeen Taqatqa, Mercedes Bentata, Shiri Jaffe-Herman, Ophir Geminder, Reyut Lewis, Marc Gotkine, Maayan Salton, Yotam Drier

## Abstract

Structural Maintenance of Chromosomes Flexible Hinge Domain Containing 1 (SMCHD1) is a non-canonical member of the structural maintenance of chromosomes (SMC) protein family involved in the regulation of chromatin structure, epigenetic regulation, and transcription. Mutations in SMCHD1 cause facioscapulohumeral muscular dystrophy type 2 (FSHD2), a rare genetic disorder characterized by progressive muscle weakness and wasting, believed to be caused by aberrant expression of DUX4 in muscle cells. Here we suggest a new role for SMCHD1 as a regulator of alternative splicing in various cell types. We demonstrate how SMCHD1 mutations cause splicing alterations of DNA Methyltransferase 3 Beta DNMT3B which can lead to hypomethylation, DUX4 expression, and FSHD pathogenesis. Analyzing RNA-seq data from muscle biopsies of FSHD2 patients and Smchd1 knocked out cells, we found that hundreds of genes were mis-spliced upon loss of SMCHD1. At least 20% of mis-spliced genes were associated with abnormalities of the musculature. Moreover, we show that mis-spliced exons tend to be bound by SMCHD1, and these exons demonstrate a slower elongation rate, suggesting SMCHD1 binding promotes exon exclusion by slowing RNA polymerase II (RNAPII). Specifically, we discovered that SMCHD1 mutations promote the splicing of the DNMT3B1 isoform of DNMT3B by perturbing RNAPII elongation rate and recruitment of the splicing factor RBM5. The mis-splicing of DNMT3B leads to hypomethylation of the D4Z4 region and DUX4 overexpression. These results suggest that mis-splicing by SMCHD1 may play a major role in FSHD2 pathogenesis by promoting the mis-splicing of different targets including DNMT3B, and highlight the potential for targeting splicing as a therapeutic strategy for this disorder.

**Significance statement:** Our study sheds light on how the loss of SMCHD1 drives the pathogenesis of facioscapulohumeral muscular dystrophy (FSHD), a rare genetic disorder characterized by muscle weakness and wasting. We found that SMCHD1 mutations led to changes in splicing of hundreds of genes, 20% of which were related to muscle abnormalities. We found that SMCHD1 tends to bind mis-spliced exons and that its binding slows down the elongation rate of RNA polymerase II often leading to the exclusion of the exon. One of these targets is DNA Methyltransferase 3 Beta (DNMT3B), and we show that the isoform promoted by SMCHD1 mutations leads to hypomethylation of a repeat region near DUX4 and to DUX4 overexpression, a known cause for FSHD. Our results provide insight into the molecular mechanisms underlying this disorder, and suggest splicing modulation as a therapeutic strategy for FSHD.

## Introduction

Splicing of precursor mRNA (pre-mRNA) is a key regulatory process in gene expression. While splicing is the process in which exons are joined together, alternative splicing is the process in which different exons are spliced together in a combinatorial fashion, either including or excluding an exon form the final transcript. Thus, alternative pre-mRNA splicing enables the construction of distinct protein isoforms from the same gene and contributes to the cell’s diverse protein population. Proper splicing is necessary for the cell’s function and when altered may lead to several pathological processes such as cancer and genetic diseases^1,2^. As an essential regulatory process in the cell, alternative splicing is tightly regulated. Many factors can regulate alternative splicing, including cis-acting elements within the pre-mRNA molecule and trans-acting factors, mostly splicing factors, proteins that alter splicing by regulating splice site selection.

Another trans-acting factor regulating alternative splicing is RNA polymerase II (RNAPII). Typically, slow RNAPII elongation kinetics promotes exon inclusion, as it exposes additional splice sites^3^. Therefore, altering the elongation rate by replacing a gene’s promoter or altering its chromatin can impact splicing^4^. In some cases, the opposite effect is observed whereby slow RNAPII kinetics promotes exon exclusion^5^. This effect was attributed to an inhibitory splicing factor that is recruited by RNAPII Carboxy-terminal domain (CTD) and gains a binding opportunity when RNAPII is slow to transcribe^5^. Several chromatin modulators have previously been shown to act as regulators of splicing, including the architectural regulator CCCTC-binding factor (CTCF), which regulates splicing via changes in RNAPII elongation rate^6,7^.

In our previous work, we conducted an unbiased high-throughput screen to identify chromatin regulators with a role in modulating alternative splicing^8,9^. Our results identified 16 chromatin proteins associated with alternative splicing regulation, including the structural maintenance of chromosomes flexible hinge domain-containing 1 (SMCHD1) protein^8^. SMCHD1 is an SMC protein comprising an N-terminal GHKL (gyrase, Hsp90, histidine kinase, MutL) ATPase domain and an SMC (structural maintenance of chromosomes) hinge domain that possesses chromatin-binding activity^10^. As an SMC protein, SMCHD1 function is speculated to contribute to the maintenance of DNA and chromatin structure. SMCHD1 knockout has previously presented significant effects on histone modifications, DNA methylation, CTCF occupancy, and chromosomal interactions^10–16^. SMCHD1 was also shown to play a key role in the X inactivation process and in the silencing of autosomal genes^10^.

A variety of heterozygous loss-of-function mutations in the SMCHD1 gene were described to cause facioscapulohumeral dystrophy type 2 (FSHD2), a late-onset progressive muscular dystrophy disease^17^. The suggested molecular basis of FSHD2 is hypomethylation of the D4Z4 macrosatellite array, caused by SMCHD1 loss of function. This results in higher expression of the DUX4 transcription factor and myocyte toxic genes^18^. FSHD1 is caused by contractions of the D4Z4 macrosatellite array, associated with loss of methylation at this site, and is more common than FSHD2^19^. While most FSHD cases that are not attributed to D4Z4 contraction are caused by SMCHD1 mutations, FSHD has also been described in patients with mutations in the DNA Methyltransferase 3 Beta (DNMT3B) gene (FSHD4) and the Ligand Dependent Nuclear Receptor Interacting Factor 1 (LRIF1) gene (FSHD3)^20,21^. Both SMCHD1 and DNMT3B variants were identified as modifiers of disease severity in FSHD1 patients as well^20,22,23^. In addition, missense mutations in the ATPase domain of SMCHD1 were demonstrated to cause Bosma arhinia microphthalmia syndrome (BAMS), a rare condition characterized by severe facial abnormalities, especially in the nasal area. The underlying molecular basis of the disease is not well understood^24–26^.

Here we describe a novel role for SMCHD1 as a regulator of alternative splicing. We studied genome-wide alternative splicing, in neural stem cells of mice with SMCHD1 mutations as well as the muscle of FSHD2 patients. We found aberrantly spliced genes to be bound by SMCHD1 on the DNA level and to be enriched for FSHD pathology. Our results show that the binding of SMCHD1 to abnormally spliced genes is associated with RNAPII pause sites. In particular, we focus on DNMT3B and demonstrate that SMCHD1 mutations lead to preferential inclusion of exons 5, 21 and 22, and expression of the full DNMT3B isoform instead of shortened DNMT3B3ΔEx5 isoform. We show that expression of the full isoform instead of DNMT3B3ΔEx5 leads to hypomethylation of the D4Z4 repeat array and promotes DUX4 expression, the key events causing FSHD. Therefore, we suggest a new model for FSHD2 pathogenesis, driven by SMCHD1-mediated alternative splicing.

## Results

### Differential splicing in FSHD2 patients with SMCHD1 mutation

To explore alternative splicing in SMCHD1 mutated FSHD2 patients, we compared RNA-seq data of muscle biopsies of four FSHD2 patients with mutations in the SMCHD1 gene with four healthy individuals^18^. While 117 genes are differentially expressed (FDR < 5%, DESeq2), at least 871 genes are mis-spliced (FDR < 5%, rMATS). 1420 mis-splicing events were identified, 46% of them are exon skipping events (649 events) (Fig. 1A, SI Appendix, Table S1. The other 54% are composed of different alternative splicing events including mutually exclusive exons (407 events), intron retention (124 events), alternative 3’ splice site (97 events), and alternative 5’ splice site (143 events). Interestingly, only eight genes were identified as both differentially expressed and alternatively spliced, suggesting that SMCHD1 independently regulates gene expression and splicing (Fig. 1B). In order to control for splicing changes that are independent of SMCHD1 mutations, we repeated the analysis comparing FSHD2 to FSHD1 patients and discovered 1968 splicing events in 845 genes (FDR < 5%, rMATS). Importantly, 253 of these genes (30%) are the same genes detected in the comparison to healthy individuals (p < 0.00001, Fisher exact test, Fig. 1C). Overall, these results reveal significant mis-splicing in SMCHD1 mutated FSHD2 patients, pointing to a role for SMCHD1 in alternative splicing.

**Figure 1:**
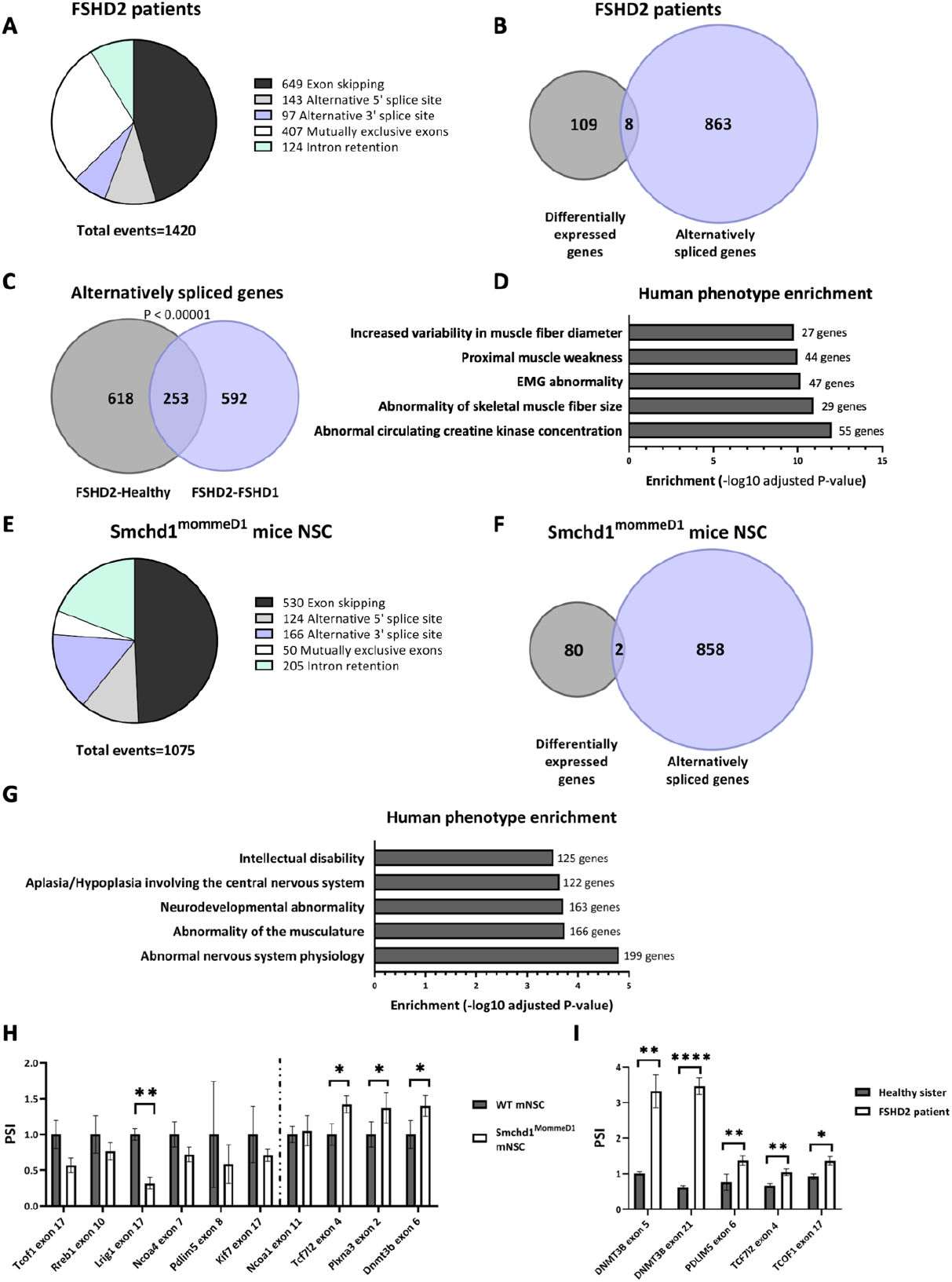
SMCHD1 is a regulator of alternative splicing. **(A-D)** Significant alternative splicing in muscles of FSHD2 patients revealed by rMATS analysis of RNA-seq data from healthy, FSHDl and FSHD2 patients (ref 18). **(A)** Proportion of each alternative splicing event between muscles of FSHD2 patients and healthy individuals. **(Bl** Venn diagram presenting the overlap between differentially expressed and alternatively spliced genes in FSHD2 patients. (C) Venn diagram presenting the overlap of alternative splicing events when FSHD2 patients are compared to FSHDl patients or to healthy individuals. (D) Top five significant events for human phenotypes gene set enrichment of FSHD2 alternatively spliced genes, bars present -log_10_ adjusted p value, number of alternatively spliced genes are annotated next to bar. **(E-H)** Significant alternative splicing in Smchdl null neural stem cells revealed by rMATS analysis of RNA-seq from three Smchdl null and two WT mice NSC samples. **(E)** Proportion of each alternative splicing event is presented relative to WT. **(F)** Venn diagram presenting the overlap between Smchdl expressed and alternatively spliced genes in Smchdl null mice NSC. **(G)** Top five significant events for human phenotypes gene set enrichment of Smchdl null alternatively spliced genes, bars present -log_10_ adjusted p value. (H) Real-time PCR was conducted to measure relevant splicing change and total mRNA amount. Results are shown as percent spliced in (PSI) calculated as exon inclusion relative to total mRNA of the gene. Values represent averages of three RNA samples relative to two control samples ±SD [*p<0.05; **p<0.01; ***p<0.001]. (I) RNA was extracted from lymphoblasts of an FSHD2 patient and her healthy sister. Real-time PCR was conducted to measure the relevant splicing event and total mRNA amount. Results are shown as percent spliced in (PSI) calculated as exon inclusion relative to total mRNA of the gene. Values represent averages of three repeats [*p<0.05; **p<0.01; ** *p<0.001;* * * * p<0.0001].

We next asked how splicing alteration by SMCHD1 can contribute to the phenotype of FSHD2. To this end, we explored the function of genes regulated in splicing by gene-set enrichment analysis, compared to all expressed (TPM>1) genes in muscle tissue. 22% (192 genes of 871, FDR < 3.2*10^−5^) of the genes mis-spliced in FSHD2 patients are associated with “abnormality of the musculature” annotation by the Human Phenotype Ontology (HPO^27^), suggesting that many genes mis-spliced due to SMCHD1 mutations in FSHD2 are associated with abnormal muscle function. The top significant enriched human phenotype terms included many annotations associated with muscular dystrophy which are related to FSHD2 pathology (Fig. 1D, SI Appendix, Table S2). Analysis of genes differentially spliced between FSHD2 and FSHD1 yielded similar results-178 genes out of 846 alternatively spliced (21%) were associated with abnormality of the musculature (FDR < 0.0002, SI Appendix, Table S2), demonstrating this is indeed due to SMCHD1 mutations and no other FSHD related mis-splicing. Finally, to further support this claim, we repeated the analysis to compare FSHD1 patients to healthy controls, and indeed in this case no significant enrichment of any muscle related phenotypes was detected. Importantly, gene-set enrichment analysis of differentially expressed genes between either FSHD2 and healthy controls or between FSHD2 and FSHD1 revealed no enrichment for FSHD-related phenotypes.

### SMCHD1 regulates alternative splicing in mice neural stem cells, embryonic fibroblasts and embryonic stem cells

To further investigate potential genes whose splicing is regulated by SMCHD1, we conducted deep sequencing of RNA from neural stem cells (NSC) sorted from Smchd1 null mice. The Smchd1^MommeD1^ mice are a previously established model with a mutation in one allele of Smchd1, resulting in Smchd1 haploinsufficiency^10^. Here, we compared MommeD1 homozygous NSCs to WT controls, to measure aberrant splicing in the absence of Smchd1. Differential splicing analysis identified 1075 splicing events in 860 genes (FDR < 5%, rMATS). 49% of the splicing events due to Smchd1 loss are exon skipping events (530 events) (Fig. 1E, SI Appendix, Table S1). The other 51% are composed of different splicing events including mutually exclusive exons (50 events), intron retention (205 events), alternative 3’ splice site (166 events), and alternative 5’ splice site (124 events). 59% of these events presented higher inclusion in Smchd1^MommeD1^ samples and 41% presented higher exclusion, suggesting Smchd1 is regulating splicing in both directions. We identified only 82 differentially expressed genes (DESeq2, FDR < 5%) and only two genes were both differentially expressed and alternatively spliced (Fig. 1F). As in FSHD2 patients, gene-set enrichment analysis for differentially spliced genes in the NSC revealed significant enrichment for phenotypes related to muscular dystrophy. 19% of mis-spliced genes (166 of 860 genes, FDR < 0.00018) are related to the “Abnormality of the musculoskeletal system” HPO (Fig. 1G, SI Appendix, Table S2).

To validate the splicing analysis, we chose ten leading candidate genes that were significantly mis-spliced and matched either an HPO annotation of “abnormal muscle physiology” or a GO annotation of “anatomical structure development”. We performed qPCR on RNA from the same cells and found four of the ten genes to be significantly mis-spliced (p < 0.05, Student’s t-test) (Fig. 1H). One of the genes that presented a change in splicing has also demonstrated a change in total expression levels. To validate the mis-splicing of the top five significant events in patients we performed qPCR in FSHD2 patient lymphoblastoid cells (SI Appendix, Fig. S1A-C), compared to a healthy sister (Fig. 1I). All five events showed the expected change in splicing, and no significant change was detected in total expression levels (SI Appendix, Fig. S1D).

To estimate the global impact of Smchd1 on alternative splicing in different cell types, we also examined mouse embryonic fibroblasts (MEF) and mouse embryonic stem cells (mESC). We reanalyzed available RNA-seq data from Smchd1^MommeD1^ homozygous mutant female MEFs^13^ and found 3997 alternative splicing events in 2349 genes (rMATS, FDR < 5%, SI Appendix, Fig. S1E, SI Appendix, Table S1). Analysis of RNA-seq data from Smchd1-KO mESC^28^ revealed a significant change in alternative splicing as well, with 1950 events in 1100 genes. (SI Appendix, Fig. S1F, Table S1). Gene-set enrichment analysis for differentially spliced genes in MEFs as well as mESC revealed significant enrichment for phenotypes related to muscular dystrophy (SI Appendix, Fig. S1G-H, Table S2). Together these findings indicate that Smchd1 regulates the alternative splicing of thousands of genes across multiple cell types, independently of its role in gene expression regulation. The identity of the alternatively spliced genes may change between cell types, but many of them are shared between two or more cell types (p < 0.00001, Fisher’s exact test, SI Appendix, Fig. S1I). Genes whose splicing is regulated by Smchd1 tend to be related to muscle dystrophy, even in unrelated cell types, demonstrating the potential of Smchd1 mediated splicing to affect muscle dystrophy, that can be unleashed in muscle cells.

## Smchd1 binding is enriched at mis-spliced exons

Next, we wanted to explore how Smchd1 regulates splicing. Since Smchd1 is a chromatin factor we wanted to explore the chromatin landscape at its splicing targets. We began by exploring whether Smchd1 directly binds mis-spliced exons. To this end, we analyzed available Smchd1-GFP ChIP-seq data from neural stem cells of wild-type (WT) mice^11^. First, we assessed Smchd1 binding in the vicinity (5kb) of mis-spliced exons and compared them to all exons expressed in either WT or Smchd1^MommeD1^ NSC, regardless of whether they are mis-spliced or not. We found Smchd1 binds 149 (13%) of mis-spliced exons, compared to 7% of all expressed exons (Fig. 2A, p < 0.0001, Fisher’s exact test). Overall, Smchd1 binds mis-spliced exons 1.92 times more often than all expressed exons. (Fig. 2A&B). Moreover, we compared Smchd1 binding in mis-spliced exons to other exons in the same genes that are not alternatively spliced, and found Smchd1 binds the mis-spliced exons 2.1 times more often. (Fig. 2B).

**Figure 2:**
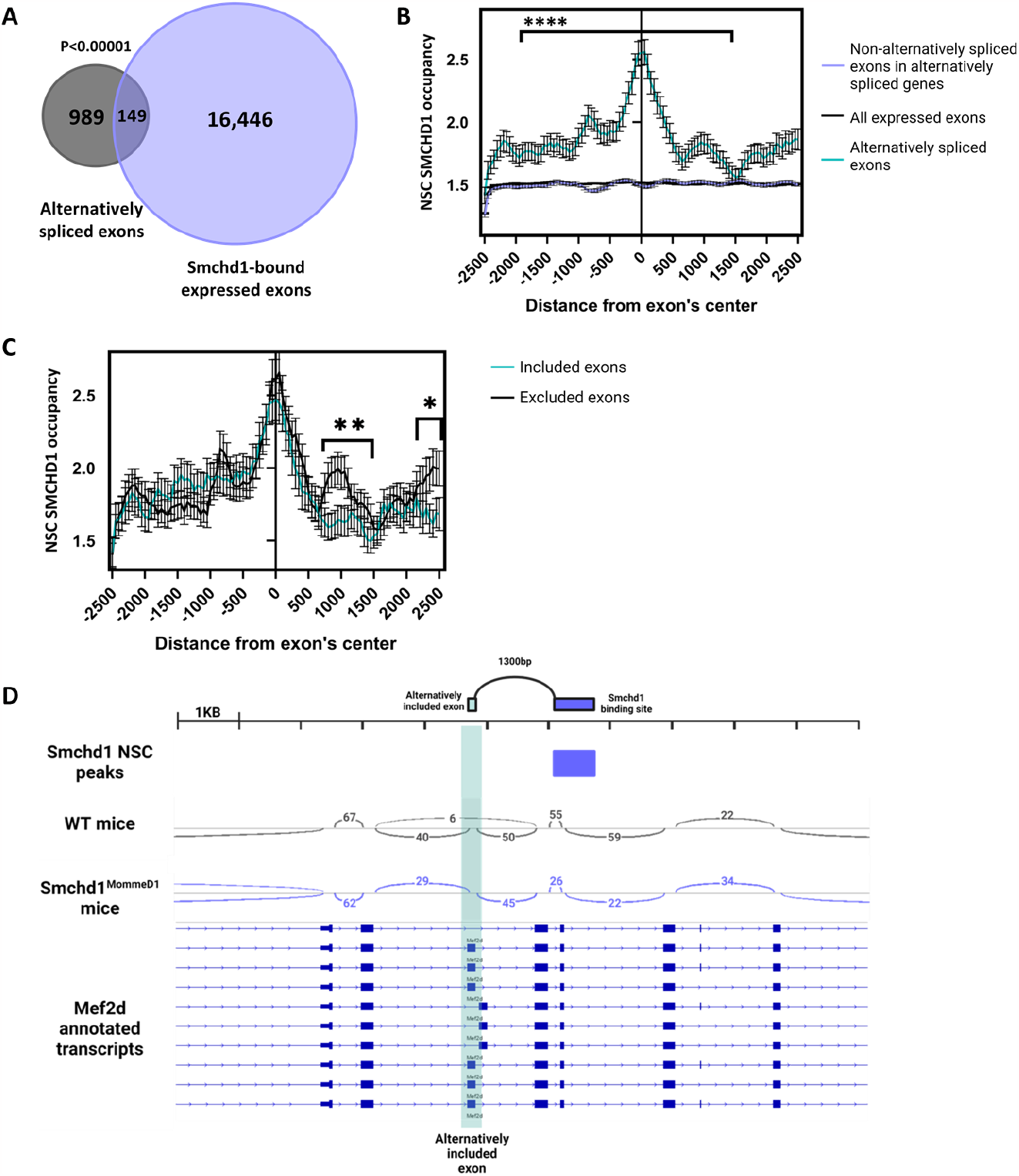
Smchdl binding is enriched downstream of its regulated excluded exons. **(A-D)** Reanalysis of GFP ChlP-seq in primary NSCs with endogenous Smchdl-GFP fusion protein, analysis was limited to expressed genes only (TPM>1). (A) Venn diagram presenting the overlap of alternatively spliced exons and exons with nearby (<5kbp) Smchdl binding site. **(B)** Aggregation plot depicting the average normalized Smchdl occupancy, at and near exons alternatively spliced (turquoise), all expressed exons (black) or non-alternatively spliced exons (purple) in alternatively spliced genes, showing stronger binding of Smchdl alternatively spliced exons [****p<0.0001]. X axis represent bins of size 50 bp around the center of the exon. **(C)** Aggregation plot depicting the average normalized Smchdl occupancy at exons differentially included or excluded in Smchdl^MommeD1^ mice [*p<0.05, **<0.01], X axis represent bins of size 50 bp around the center of the exon. **(D)** Genome browser view of the Mef2d alternatively spliced junctions presented by sashimi plots, arcs denote splice junctions quantified in spanning reads. Mutually exclusive alternatively spliced exon is highlighted in turquoise. Refseq transcripts are presented as a reference.

Interestingly, we found that Smchd1 preferentially binds downstream to exons that are included in Smchd1 null cells, but excluded when bound by Smchd1 in WT cells (“excluded exons”) (p < 0.01, Wilcoxon test) (Fig. 2C). For example, exon 5 of Mef2d, a member of the myocyte-specific enhancer factor 2 (Mef2) family involved in muscle cell differentiation and development^29^. Exon 5 is adjacent to an Smchd1 binding site and it is skipped in WT mice but included in Smchd1^MommeD1^ mice (Fig. 2D). We repeated the analysis for another available NSC Smchd1-GFP ChIP-seq dataset^30^ and found Smchd1 binds 1.38 more to mis-spliced exons compared to all expressed exons (p < 0.00001, Fisher’s exact test) and specifically binds more to excluded exons (p < 0.009, student’s t-test, SI Appendix, Fig. S2A-C). Moreover, we repeated this analysis for MEFs, utilizing published Smchd1 binding data in MEF^13^. We observed a similar association with splicing: Mis-spliced exons had 16% more Smchd1 binding sites than all expressed exons (p < 0.044, Fisher’s exact test), and Smchd1 binding preferentially binds downstream of excluded exons (SI Appendix, Fig. S2D). Additionally, we repeated the analysis for a 1 kb window around the exon and found similar enrichment of Smchd1 in all datasets. Smchd1 binds 4-times and 5.5-times more to alternatively spliced exons compared to all expressed exons in the NSC datasets (P < 0.00001, Fisher’s exact test), and 3 times more (P < 0.00001, Fisher’s exact test) in MEFs. This significant overlap between Smchd1 binding and alternatively spliced exons suggest direct effect of Smchd1 binding on alternative splicing. The relatively modest effect sizes suggests that some genes may be also indirectly affected, and some Smchd1 binding sites may not yield a detectible difference in splicing. Overall, our analysis demonstrates that the binding of Smchd1 is enriched downstream of excluded exons, suggesting Smchd1 loss leads to aberrant inclusion of the exons.

## Smchd1 binding correlates with RNAPII stalling at mis-spliced exons

Since RNAPII elongation rate can regulate alternative splicing we next asked whether Smchd1 binding is associated with RNAPII elongation. To address this, we reanalyzed ChIP-seq data of phosphoserine 2 of RNAPII carboxy-terminal domain (pSer2), a marker of elongating RNAPII, from C2C12 mouse myoblasts^31^. Enrichment of pSer2 marks a slow elongation rate or stalling of RNAPII^32^. We compared the RNAPII pSer2 signal between mis-spliced exons with and without Smchd1 binding. Our analysis revealed a significant enrichment of RNAPII pSer2 in the exons bound by Smchd1 (Fig. 3A). We repeated the analysis for Smchd1 binding sites identified in a different NSC dataset^30^ or MEFs^13^ and also found a significant enrichment of RNAPII pSer2 at Smchd1-bound mis-spliced exons (SI Appendix, Fig. S3A-B). Moreover, we compared RNAPII pSer2 signal between included or excluded exons and found a specific enrichment at excluded exons (Fig. 3B). Together, these results suggest that Smchd1 binding in mis-spliced exons is correlated with RNAPII stalling and exon exclusion.

**Figure 3:**
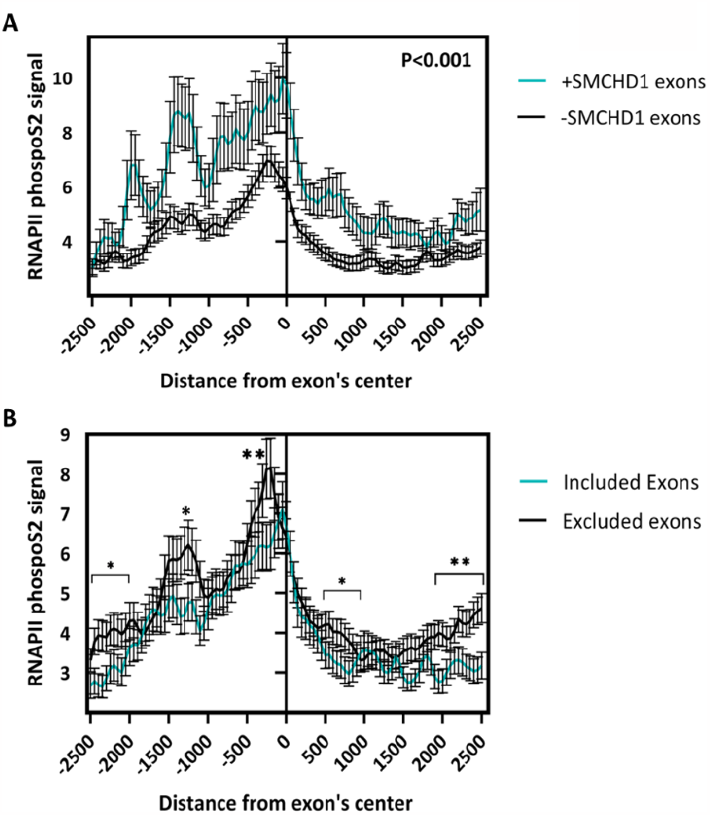
Smchd1 binding is associated with slow elongation 2ra1t4e of RNAPII. **(A-B)** Aggregation plots depicting the average normalized phospho-Ser2 levels of RNAPII at and near alternatively spliced exons differentially bound by Smchd1 **(A)** and at and near exons differentially included or excluded in Smchd1^MommeD1^ mice **(B)** in C2C12 cells. X axis represent bins of size 50 bp around the ce2nt1e7r of the exon [* FDR<0.05; ** FDR<0.01; *** FDR<0.001; ****p<0.0001].

### DNMT3B splicing is regulated by SMCHD1 in mice and human

Our analysis revealed that DNA methyltransferase 3 beta (DNMT3B) splicing is regulated by SMCHD1 in both human FSHD2 patient (Fig. 1I), mice NSCs (Fig. 1H, SI Appendix, Fig. S4A), MEFs (rMATS FDR < 0.0004) and mice ESC (rMATS FDR < 0.003). DNMT3B presented significant splicing changes in exons 5 and 21-22 in human, corresponding to exons 6 and 20-21 in mice. DNMT3B is known to have dozens of alternatively spliced isoforms with distinct functions^33^. DNMT3B exons 21 and 22 encode part of the MTAse catalytic domain of the protein while the exon 5 region does not contain any known functional domains. DNMT3B3ΔEx5 isoform is known to be associated with increased DNA binding affinity and enhanced cell growth^33^. Mutations of DNMT3B in FSHD are associated with D4Z4 hypomethylation and with high levels of DUX4 expression^20^. Specifically, a mutation in DNMT3B MTAse catalytic domain was shown to cause FSHD^20^. Alternative splicing of DNMT3B may affect D4Z4 methylation and thus contribute to disease development. To validate SMCHD1-mediated alternative splicing of DNMT3B in human cells, we knocked down SMCHD1 by siRNA in HCT116 cells (SI Appendix, Fig. S4B), a cancer cell line with expressed and active DNMT3B. qPCR analysis showed a 60% increase in the inclusion of exon 5 (p < 0.05, student t-test) and a two-fold increase in the inclusion of exon 21 (p < 0.01, student t-test) (Fig. 4A-B). Semi-quantitative PCR analysis showed a three-fold increase in exon 5 inclusion and an 18-fold increase in exons 21-22 inclusion upon SMCHD1 knock-down (Fig. 4C-D, SI Appendix, Fig. S4C-D). Overall, this demonstrates DNMT3B alternative splicing is indeed regulated by SMCHD1.

**Figure 4:**
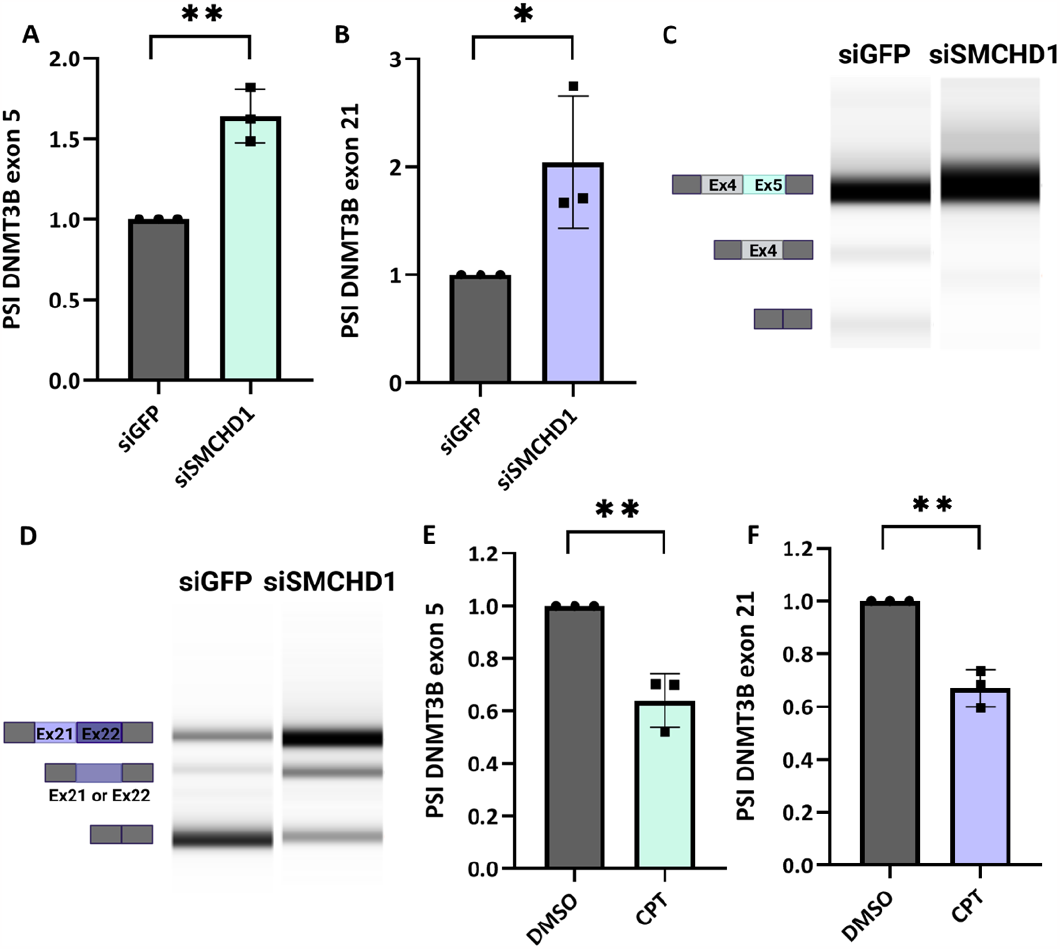
DNMT3B exon 5 and 21 are regulated by SMCHD1 and RNAPII stalling. **(A-D)** HCT116 cells were transfected with siRNA targeting SMCHD1 and GFP as negative control. Total RNA was extracted and analyzed by real-time PCR for DNMT3B exon 5 **(A)** and exon 21 **(B)** inclusion relative to DNMT3B total mRNA amount. PSI was calculated by dividing exon inclusion in DNMT3B total mRNA amount. Semi quantitative PCR was conducted for exons 4-5 **(C)** and exons 21-22 **(D). (E-F)** HCT116 cells were treated with 6uM CPT or DMSO as negative control, for 6 hr. Total RNA was extracted and analyzed by real-time PCR for DNMT3B exon 5 **(E)** and exon 21 **(F)** inclusion relative to DNMT3B total mRNA amount. PSI was calculated by dividing exon inclusion in DNMT3B total mRNA amount. Values represent averages of three independent experiments ± SD; [* p<0.05; **p<0.01] (paired Student’s t-test).

Our previous results suggested that SMCHD1 regulates alternative splicing by binding to the vicinity of the alternative exon and slowing RNAPII elongation rate. To investigate whether DNMT3B alternative splicing is mediated by slower RNAPII, we treated HCT116 cells with CPT, a topoisomerase inhibitor that slows RNAPII elongation. We used a low dose of 6uM CPT to slow down RNAPII without stopping it, while also avoiding any potential DNA damage. Our results show a 70% reduction in total mRNA amount of DNMT3B and a 35% reduction in exon 5 and 21 inclusion following treatment with CPT (p < 0.01, student t-test) (Fig. 4E-F and SI Appendix, Fig. S4E), suggesting that RNAPII stalling indeed promotes exclusion of SMCHD1 regulated exons at DNMT3B gene.

### Identification of splicing factors regulating the alternative splicing of DNMT3B

Our previous findings suggest that SMCHD1 is promoting exon exclusion by RNAPII stalling. We hypothesize that RNAPII stalling by SMCHD1 promotes exclusion by recruiting splicing factors. To predict which splicing factors may be involved we performed RNA binding proteins motif analysis on exons excluded by SMCHD1. To this end, we compared excluded exons and their flanking 10 kb sequences to those of included exons in mice NSC and human FSHD2 patients. Our analysis found 35 significantly enriched motifs in human and 17 in mice. Five motifs were found enriched in both datasets: RBFOX1, CPO, MSI, RBM5, and SRSF10 (Fig. 5A).

**Figure 5:**
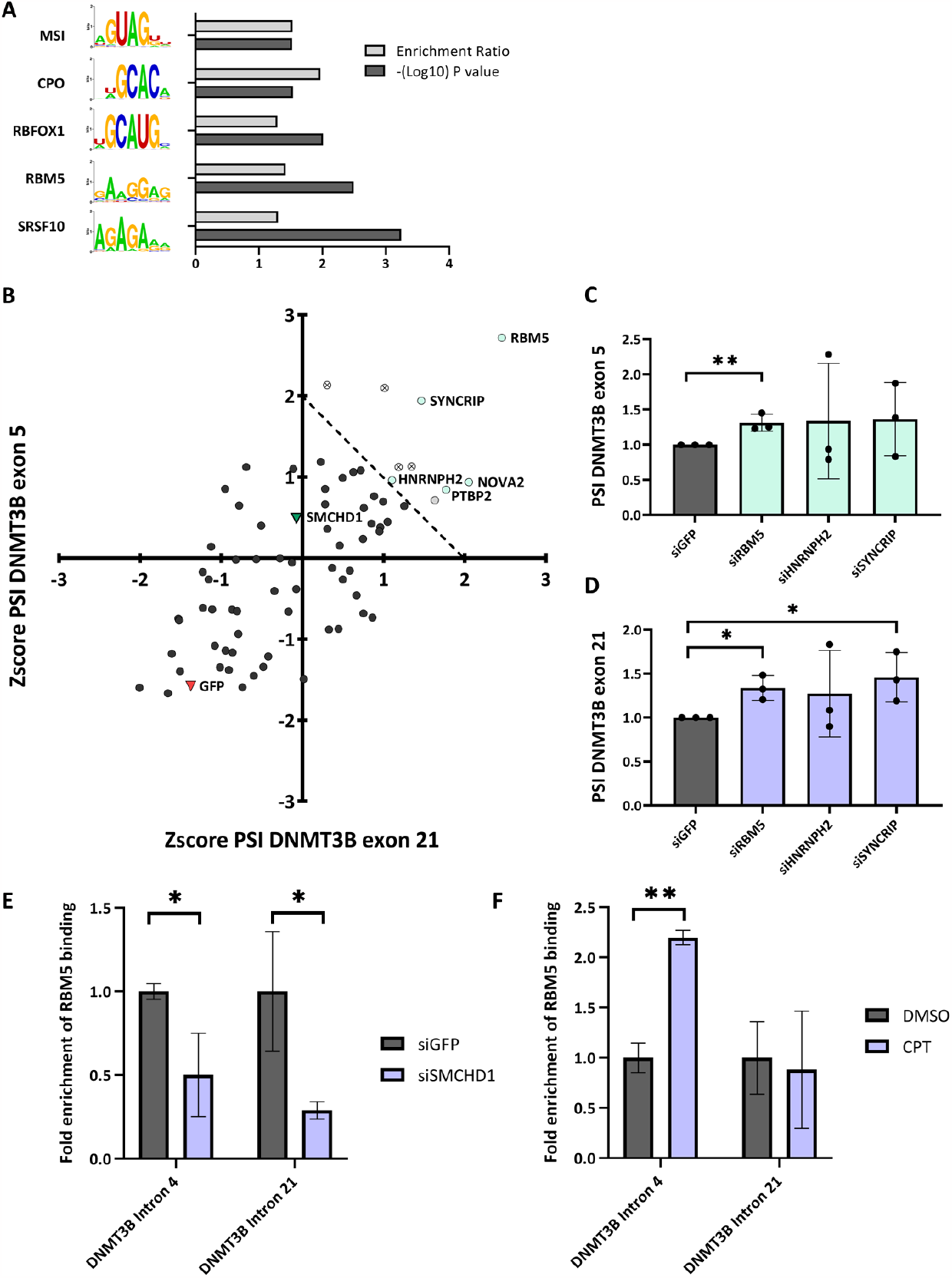
DNMT38 exon 5 and 21 are regulated by RBM5. **(A)** Enrichment of binding sites of SMCHDl regulated exons: Enriched RNA motifs in alternatively excluded exons in both mice and human compared to included exons. **(Bl** HCT116 cells were transfected with siRNA targeting 71 human splicing factor, SMCHDl as a positive control and GFP as negative control. Total RNA was extracted and analyzed by real-time PCR for DNMT3B exon 5 and exon 21. Z scores were calculated for DNMT3B exon 5 and exon 21 PSI, as DNMT3B exon inclusion/DNMT3B total mRNA and normalized to siGFP as a negative control. Shown in scatter plot: red triangle represents GFP (negative control), green triangle represents SMCHDl (positive control), gray circle represents a lowly expressed factor, crossed circles represent hits that are inconsistent between repeats and blue circles represent splicing factor hits. **(C-D)** HCT116 cells were transfected with siRNA targeting the splicing factor hits and GFP as a negative control. Total RNA was extracted and analyzed by real-time PCR for DNMT3B exon 5 (C) and exon 21 **(D)** inclusion relative to DNMT3B total mRNA amount. PSI was calculated as DNMT3B exon inclusion/DNMT3B total mRNA and normalized to negative control. Values represent averages of three repeats ±SD. Negative control (siGFP) PSI is represented by the dotted line at 1; [* p<0.05] (Student’s I-test). **(El** RNA-lmmunoprecipitation of RBMS in HCT116 transfected with siSMCHDl or negative control (siGFP) for 72 h. Real-time PCR for DNMT3B intron 4 and 21 relative to input. **(Fl** HCT116 cells were treated with GuM CPT or DMSO as negative control, for 6 hr. Real-time PCR for DNMT3B intron 4 and 21 relative to input. (E-F) Values represent averages of three technical replicates ±SD; [* p<0.05, ‥ p<0.01] (paired Student’s t-test).

To experimentally identify potential splicing factors cooperating with SMCHD1, we performed an unbiased siRNA screen with the use of DNMT3B alternative splicing as readout. Specifically, we used a library of siRNA oligos directed to the 71 human splicing factors (as described in SpliceAid-F^34^) in HCT116 cells. We monitored alternative splicing of DNMT3B exons 5 and 21 using qPCR. As SMCHD1 is regulating the alternative splicing of both exon 5 and 21 of DNMT3B, we expected that a splicing factor working with SMCHD1 will regulate both events. Z scores were calculated for the average percent spliced in (PSI) of exons 5 or 21 separately. Splicing factors for which both z scores are higher than the positive control (siSMCHD1), and for which the sum of both z scores was higher than 2 were considered for downstream analysis. The screen identified six factors: RBM5, SYNCRIP, HNRNPH2, NOVA2, PTBP2, and ELAVL3 (Fig. 5B). Of those six factors we filtered out factors with expression level below detection rate and eventually found five hits: RBM5, SYNCRIP, HNRNPH2, NOVA2 and PTBP2 (SI Appendix, Fig. S4F). To validate these hits, we conducted a secondary screen knocking down each splicing factor by siRNA in HCT116 cells and tested the inclusion of DNMT3B exons while monitoring the knock-down level of each splicing factor (SI Appendix, Fig. S4G). The silencing of PTBP2 and NOVA2 was unsuccessful in the secondary screen, and we can assume that it was similarly unsuccessful in the original screen. Therefore, we excluded them from further analysis. qPCR analysis showed a significant increase in the inclusion of DNMT3B exons 5 and 21 only for RBM5 (31% and 34% increase respectively, p<0.005 and p<0.017, student t-test) (Fig. 5C-D). Overall, motif enrichment and splicing factor screen suggest RBM5 as a potential ally of SMCHD1 in its alternative splicing regulation.

### DNMT3B alternative splicing is regulated by SMCHD1 via RBM5

We hypothesize that SMCHD1 binding to the DNA stalls RNAPII to allow RBM5 binding to pre-mRNA. To check our hypothesis, we silenced SMCHD1 and measured the binding of RBM5 using RNA-IP to DNMT3B pre-mRNA. To this end, we performed RNA-IP for RBM5 in HCT116 cells. Monitoring DNMT3B introns 4 and 21 relative to input, we found silencing of SMCHD1 led to a reduction of 50% and 70% in RBM5 binding, respectively (p<0.013 and p<0.017, student t-test) (Fig. 5E). This result suggests that SMCHD1 is a regulator of RBM5 binding to DNMT3B’s pre-mRNA.

To test whether RNAPII stalling mediates RBM5 binding we performed RNA-IP following CPT treatment and found a 2-fold increase in RBM5 binding to DNMT3B intron 4 (p<0.004, student t-test) while there was no change in RBM5 binding to DNMT3B intron 21 (Fig. 5F). Overall, these results support the hypothesis that the RBM5 and SMCHD1 proteins functionally interact in the regulation of DNMT3B alternative splicing and that RBM5 recruitment to DNMT3B exon 5 is affected by RNAPII stalling.

### DNMT3B alternative splicing promotes D4Z4 hypomethylation and DUX4 expression

SMCHD1 mutations promote aberrant inclusion of both exons 5 and 21-22 of DNMT3B, and DNMT3B mutations were previously associated with DUX4 expression and D4Z4 hypomethylation^20^. Thus, we next aimed to test whether DNMT3B isoforms cause hypomethylation of the D4Z4 region and promote DUX4 expression. To test the impact of each DNMT3B isoform on D4Z4 methylation, we infected DNMT3B-null HCT116 cells with either GFP-DNMT3B1, GFP-DNMT3B3ΔEx5 or an empty GFP vector (Fig. 6A-C, SI Appendix, Fig. S5). We determined DNA methylation levels after 14 days at three regions of the D4Z4 array using bisulfite-PCR sequencing and found a significant decrease in D4Z4 methylation in DNMT3B1 expressing cells compared to DNMT3B3ΔEx5 cells (p < 0.00001, student’s t-test) (Fig. 6D). Interestingly, DNMT3B1 cells presented decreased methylation level in several regions compared to cells infected with empty GFP vector, and therefore no active DNMT3B at all (p < 0.02, student’s t-test) (SI Appendix, Fig. S5). Overall, these results show that DNMT3B alternatively spliced isoforms differentially regulate DNA methylation, and specifically at the D4Z4 locus. Next, we assessed the expression level of DUX4 mRNA in these cell lines by PCR and found that while DNMT3B3ΔEx5 expressing cells presented only a minimal expression of DUX4, DNMT3B1 expressing cells show significantly higher levels (Fig. 6E, SI Appendix, Fig. S5B). Together with our finding that SMCHD1 mutations shift DNMT3B splicing towards the DNMT3B1 isoform, this suggests that SMCHD1 mutations drive FSHD2 pathogenesis by altering DNMT3B splicing and prompting D4Z4 hypomethylation and DUX4 expression.

**Figure 6:**
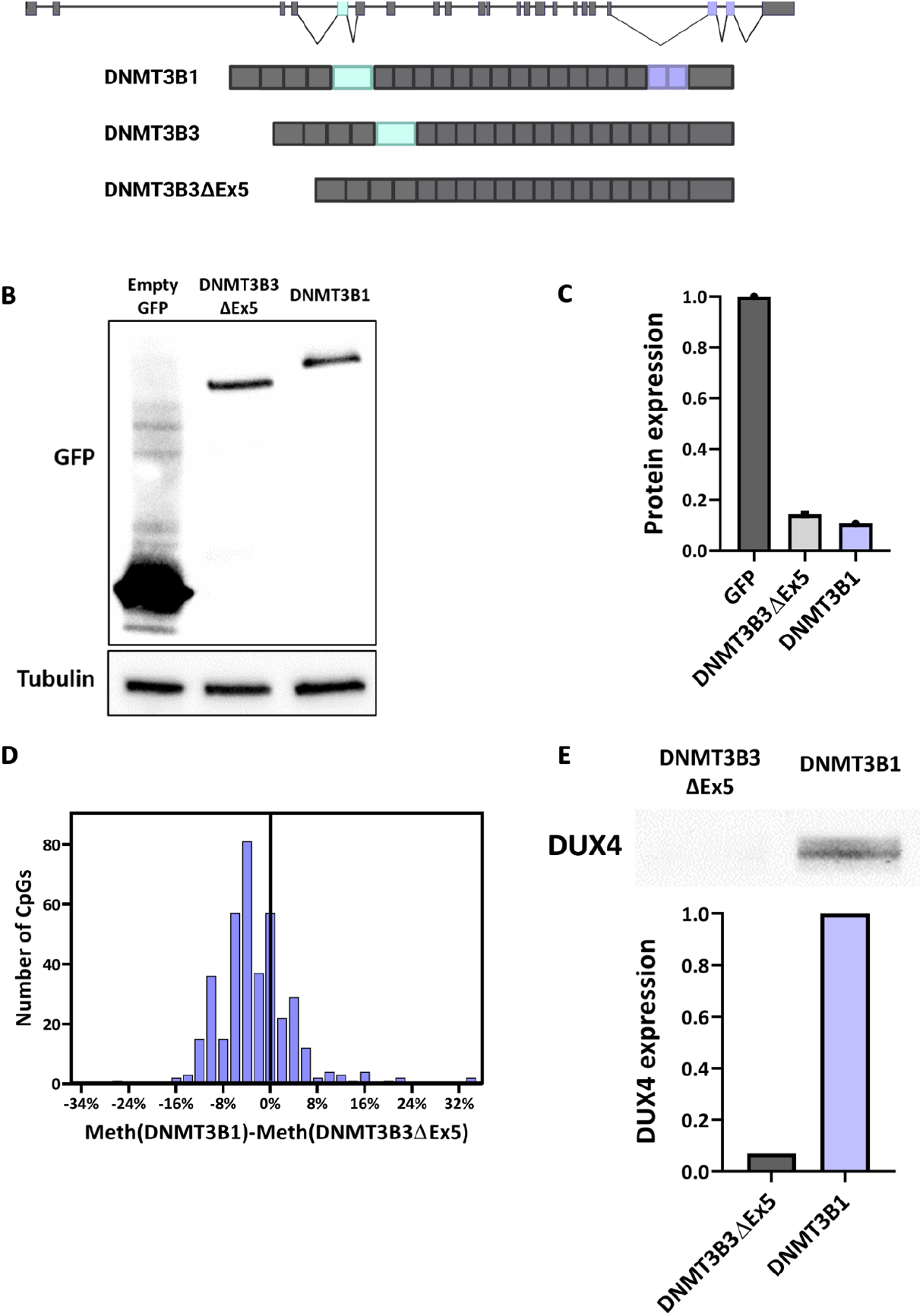
DNMT3B1 isoform reduces methy/ation at the D4Z4 region leading to increase of DUX4 expression. **(A)** Schematic representation of the *DNMT38* gene and its alternatively spliced isoforms. **(B-C)** DNMT3B null HCT116 cells were infected with lentiviruses containing empty-GFP, GFP-DNMT3B3fiEx5 or GFP-DNMT3Bl. Western blot conducted with the indicated antibodies **(B)** shows successful infection. Bands were quantified and normalized to Tubulin (C). **(D)** DNA was isolated from cells with each of the isoforms and converted with bisulfite. PCR products of three locations along the D424 region were sequenced, and methylation levels in each CpG were assessed using Biscuit. Histogram presents difference in methylation level measured over 470 CpGs in the D424 region. **(E)** RNA was extracted from HCT116 cells with the DNMT3B3tiEx5 and DNMT3Bl isoforms. Semi quantitative PCR was conducted for DUX4 mRNA level and bands were quantified.

## Discussion

In this work, we identify a novel role for SMCHD1 as a splicing regulator and suggest a mechanism for its action. While SMCHD1 is known to regulate chromosomal interactions and gene expression, here we show for the first time that it is also a splicing regulator. While previous reports did not detect differential isoform usage in Smchd1^MommeD135^, here we profiled mRNA by very deep sequencing accompanied by detailed differential spicing analysis allowing a more robust detection of exon inclusion and exclusion events. We suggest a mechanism in which SMCHD1 binding modulates RNAPII elongation rate. We hypothesize that RNAPII stalling is allowing the recruitment of inhibitory splicing factors and identify RBM5 as a potential ally in SMCHD1 regulatory pathway (Fig. 7).

**Figure 7:**
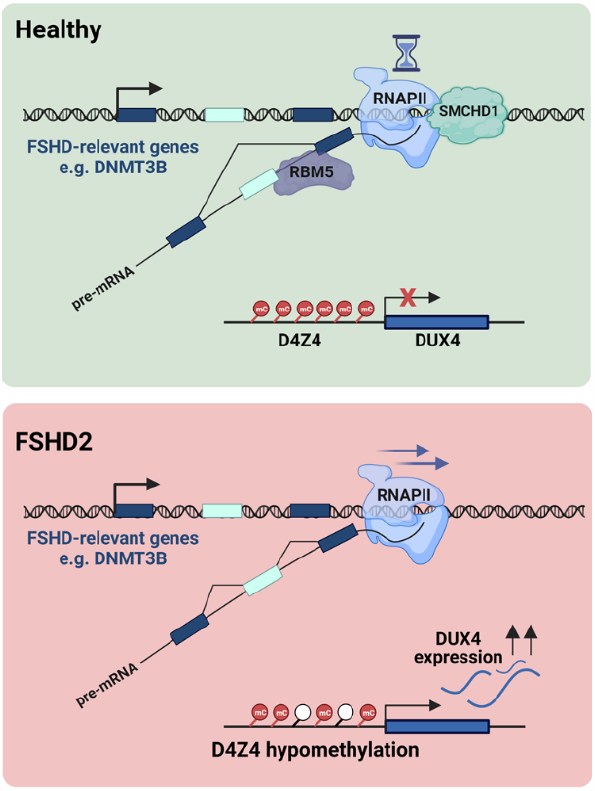
Model for SMCHD1 pathophysiology in FSHD2 driven by its abnormal splicing. In healthy cells, SMCHD1 binding is specifically enriched in the proximity of alternative exons, particularly excluded exons. A slower rate of RNAPII elongation is linked to SMCHD1 binding and related to exon exclusion. Excluded exons are characterized by a high density of RBM5 motifs, which inhibit exon inclusion and promote exon exclusion. However, in FSHD2 muscle cells, SMCHD1 mutations lead to abnormal exon inclusion. This mis-splicing of FSHD-related genes, including DNMT3B, results in decreased methylation of the D4Z4 region and increased expression of the DUX4 gene.

Several chromatin modulators have previously been shown to act as regulators of splicing, and different mechanisms for chromatin-mediated splicing regulation were identified. Our results suggest that SMCHD1 regulation of splicing is mediated by RNAPII kinetics, similar to the mechanism suggested for CTCF-mediated splicing regulation^6,7^. While CTCF is slowing RNAPII to promote exon inclusion, we found that SMCHD1-mediated slow RNAPII kinetics is correlating with exon exclusion (Fig. 3B) and is mediated by recruitment of splicing factors and specifically RBM5 (Fig. 5A-D). Specifically, we show that SMCHD1 binds the vicinity of its target exons to stall RNAPII and allow for RBM5 binding to pre-mRNA (Fig. 2B, 3A, 5E-F). Overall, our results reveal a new mechanism for chromatin-mediated splicing regulation.

We found SMCHD1 to regulate the splicing of multiple genes with a potential clinical significance for FSHD2 development. We found aberrant splicing of hundreds of genes that are known as key factors for myocyte function and mutations in dozens of them were previously identified as pathogenic in muscular dystrophy. For example, Titin (TTN) is mis-spliced in FSHD2 patients. Mutations in TTN are a known cause for several muscular dystrophies^36^ including limb-girdle muscular dystrophy (LGMD) and has many alternatively spliced isoforms with distinct functions^37^, suggesting that splicing alteration of this gene can be associated with muscular pathology. Another example is the *calpain 3 (CAPN3)* gene which is mis-spliced in FSHD2 patients and its mutations are causing LGMD as well^38^. *CAPN3* was previously identified as significantly alternatively spliced in FSHD1 patients and its alternative splicing resulted in muscle cells differentiation defect^39^, suggesting it has different functioning isoforms. We found that *CAPN3* is also differentially spliced between FSHD1 and FSHD2 patients, at different exons from the previously reported one in FSHD1 (SI Appendix, Table S1). Specifically, the splicing alteration we identified in exons 15-16 was previously described to disrupt skeletal muscle mass^40^. Moreover, we found significant alternative splicing changes in *Troponin T 1 (TNNT1)* and *TNNT3* genes in FSHD2 patients. Both genes are key factors in myocyte function and have known alternatively spliced isoforms^41^.TNNT3 was previously shown as alternatively spliced in FSHD and its aberrant splicing was found to characterize dystrophic muscles in FSHD patients^42,43^ Overall, the cumulative effect of these alterations and others we identified may contribute to the phenotype of FSHD.

In this work, we focus on SMCHD1’s prominent target, DNMT3B, which is mutated in FSHD4 patients. Previous findings indicate Dnmt3b is binding to the D4Z4 array and its knockdown results in elevated DUX4 expression in several cell types^44^. While SMCHD1 mutations are known to be present in FSHD2 patients and are associated with D4Z4 hypomethylation and DUX4 expression, the specific mechanism for SMCHD1-associated hypomethylation is unclear. Our findings suggest that mutations in SMCHD1 lead to the mis-splicing of multiple genes including DNMT3B, supporting the DNMT3B1 isoform over the DNMT3B3 isoform (Fig. 1I). Finally, DNMT3B mis-splicing cause aberrant DNA methylation in HCT116 cells at D4Z4 which causes an increase in DUX4 expression (Fig. 6D-E, 7). The detected change in DNA methylation at D4Z4 is modest, yet sufficient to account for the difference in expression, likely representing a change in a subset of the cells, and therefore a modest average methylation change. DNMT3B mediated regulation of DNA methylation occurs mostly during differentiation, and therefore loss of SMCHD1 is likely detrimental at these stages. We focused our analysis on HCT116 cells, a colon cancer stem-like cell line, where DNMT3B was shown to be expressed, active, and to play a role in DNA de novo methylation and maintenance^45^, to represent partially differentiated cells. Together, our results suggest a novel mechanism for FSHD pathogenesis orchestrated by SMCHD1 (Fig. 7).

In our work, we present that the shift from the DNMT3B3ΔEx5 isoform which does not contain the MTAse domain to the DNMT3B1 isoform which does is associated with D4Z4 hypomethylation in HCT116 cells. While the known role of the ’active’ isoform is opposing the observed finding^46^, we hypothesize that DNMT3B active-inactive isoform balance is important for keeping D4Z4 methylated by complexes involving DNMT3B. DNMT3B inactive isoforms were previously shown to enhance DNA methylation and were suggested as accessory proteins to recruit and positively regulate DNMT3A^47,48^. Specifically, previous studies indicate a gain in Dnmt3a catalytic efficiency following the presence of Dnmt3b inactive isoforms and that Dnmt3a and Dnmt3b3 form a stable complex^48^. Moreover, the reintroduction of DNMT3B inactive isoform constructs in DNMT3B null HCT116 cells was shown to cause genome-wide methylation restoration in a similar matter to that of DNMT3B1 active isoform^47^. Thus, it may be possible that the absence of the inactive isoform, which plays a crucial co-factor in DNMT3 complex activity, leads to hypomethylation. Thus, mis-splicing and downregulation of the inactive isoform may disrupt the methylation of D4Z4 and cause FSHD.

## Methods

### Cell lines

HCT116 (ATCC Number: CCL-247) cells were grown in Dulbecco’s modified Eagle’s medium (DMEM) supplemented with 10% fetal bovine serum. Lymphoblastoid cell lines were grown in RPMI-1640 supplemented with 20% fetal bovine serum. Cell lines were maintained at 37°C and 5% CO2 atmosphere.

### Ethics Statement

Human samples were obtained under the Hadassah Institutional Helsinki committee, approval no. 0198-11 HMO. All patients gave written informed consent.

### RNA sequencing and analysis

RNA from WT and Smchd1-null (Smchd1^MommeD1/MommeD1^) NSCs from embryonic day 14.5 (E14.5) mouse brains was obtained as previously described^35^. We performed next-generation sequencing, using the RNA ScreenTape kit (catalog #5067-5576; Agilent Technologies), D1000 ScreenTape kit (catalog #5067-5582; Agilent Technologies), Qubit RNA HS Assay kit (catalog # Q32852; Invitrogen), and Qubit DNA HS Assay kit (catalog #32854; Invitrogen) were used for each specific step for quality control and quantity determination of RNA and library preparation. For mRNA library preparation: TruSeq RNA Library Prep Kit v2 was used (Illumina). In brief, 1 μg was used for the library construction; the library was eluted in 20 μL of elution buffer. Libraries were adjusted to 10 mM, and then 10 μL (50%) from each sample was collected and pooled in one tube. Multiplex sample pool (1.5 pM including PhiX 1.5%) was loaded in NextSeq 500/550 High Output v2 kit (75 cycles cartridge and 150 cycles cartridge; Illumina). Run conditions were in paired-end (43 × 43 bp and 80 × 80 bp, respectively) and loaded on NextSeq 500 System machine (Illumina). NSC RNA-seq data were deposited in NCBI’s Gene Expression Omnibus and are accessible through GEO Series accession number GSE223039. FSHD patients’ RNA-seq data were obtained from the GEO database, accession GSE56787^18^. SMCHD1 mutant MEF RNA-seq data were obtained from GEO, accession GSE121184^13^. SMCHD1 KO mESC RNA-seq data were obtained from GEO, accession GSE126467^28^.

Reads were aligned to the mm10 (NSC and MEF) or hg38 (FSHD patients) using STAR version 2.7.10^49^ with default parameters. We counted reads in genes with featureCounts version 2.0.1^50^ using GENCODE release M21 (NSCs and MEFs) or GENCODE release 33 (FSHD patients). DESeq2^51^ was used to identify differentially expressed genes. rMATS (version 4.1.1)^52^ was used to identify differential alternative splicing events. For each alternative splicing event, we used the calculation on both the reads mapped to the splice junctions and the reads mapped to the exon body (JCEC). Gene-set enrichment analysis was performed using g:Profiler^53^ online tool or the R package “gprofiler2” using the HPO^27^ June 2022 release. To limit bias due to the inability to call differential splicing in lowly expressed genes, we limited the analysis only to genes with expression > 1 TPM (SI Appendix, Table S3), comparing alternatively spliced genes with > 1 TPM to all genes with > 1 TPM. The false discovery rate (FDR) was controlled by the Benjamini-Hochberg procedure.

### ChIP sequencing and analysis

Smchd1-GFP NSC ChIP-seq data were obtained from the GEO database, accession GSE111722^11,^ and GSE174066^30^. Smchd1 MEF ChIP-seq data were obtained from the GEO database, accession GSE111820^13^. RNAPII phospoS2 in C2C12 cells was obtained from ENCODE^31^, accession ENCSR000AIU. For Smchd1 ChIP, FASTQ files were obtained using SRA-toolkit (version 3.0.0) and aligned to mm10 genome with BWA-mem (version 0.7.17)^54^ using default parameters. PCR duplicates were marked and removed using Samtools (version 1.15.1)^55^.

Peak calling and annotation were performed using HOMER version 4.11^56^. Peak calling was done for Smchd1 peaks using the ’histone’ mode, otherwise default parameters were used. Peak calling was done relative to the background signal, using input for Smchd1-MEF, whole cell extract for Smchd1-GFP NSC (2021 dataset, GSE174066), and GFP ChIP-seq in WT cells for Smchd1-GFP NSC (2018 dataset, GSE111722). ChIP-seq signal estimation and visualization were done using HOMER annotatePeaks tool histogram mode, with the following parameters: -size 5000 -hist 50 -ghist. The average ChIP signal and SEM were calculated for each 50bp genomic bin. The significance of differential binding was called using the Wilcoxon test with FDR correction by the Benjamini-Hochberg procedure.

### siRNA interference

Human HCT116 cells were seeded in 6-well culture plates (1.75 × 10^5^ cells/well). After 24 hours, cells were transfected with 20 μM of esiRNA (Sigma). using TransITx2 system Mirus Bio™ (MIR2700, Thermo Fisher Scientific) following the manufacturer’s instructions. As a control, cells were transfected with esiRNA directed for GFP (EHUEGFP, Sigma). For each condition cells were seeded as triplicates and collected for examination after 72 hours.

### CPT treatment

To impede the dynamics of transcribing RNAPII, HCT116 cells were plated to achieve 60% confluency and treated with camptothecin (CPT, Sigma) to a final concentration of 6 μM for 6 h. As a control, cells were treated with DMSO at the same concentration.

### qRT-PCR

RNA was isolated using the GENEzol™ TriRNA Pure Kit (Geneaid). With the qScript cDNA Synthesis Kit (Quantabio), cDNA was synthesized from 1 μg RNA in a 20-μl reaction volume and afterward diluted to 4 ng/μl. For quantitative real-time PCR 20 ng cDNA and 1 pmol/μl primers were mixed with 2x SYBR (BioRad) in a total volume of 13 μl for each well. Cyclophilin A was used as a reference gene. Reactions were performed for 40 cycles with a Tm of 60 °C. Primers used in this study are provided in SI Appendix, Table S4.

### Semi-quantitative PCR

RNA was isolated using the GENEzol™ TriRNA Pure Kit (Geneaid). With the qScript cDNA Synthesis Kit (Quantabio), cDNA was synthesized from 1 μg RNA in a 20-μl reaction volume and afterward diluted to 4 ng/μl. The PCR reaction was done with Taq Mix Red PCR MasterMix (Tamar), 40 ng of cDNA, and 1 pmol/μl of primers.

### siRNA splicing factor screen

EsiRNA library targeting 71 human splicing factors was purchased from Sigma. HCT116 cells were seeded in a 96-well plate and reverse transfected with 50 nM of esiRNA using the Mirus TransITx2 system. We used siSMCHD1 as the positive control and siGFP (Sigma) as a negative control. After 72 hours, cells were lysed with BioRad’s SingleShot buffer. The cell lysate was used directly for cDNA synthesis followed by qPCR.

### Lymphoblastoid cell line establishment

Venous blood was collected in EDTA-coated tubes. PBMC separation was performed using Lymphoprep (Stemcell Technologies). Cells were incubated with B95 cell line media containing EBV and Cyclosporine was added for seven days of incubation.

### RNA Immunoprecipitation

Cells were washed with ice-cold PBS, harvested, and lysed for 30 min on ice in a buffer containing 0.5% NP-40 150 mM NaCl, 50 mM Tris-HCl pH = 7.5, and 1 mM EDTA supplemented with protease and RNasin inhibitors followed by sonication in an ultrasonic bath (Qsonica, Q800R2 Sonicator) for six cycles of 5 sec ON and 30 sec OFF. Supernatants were collected after centrifugation at 16,000g for 20 min. Antibodies with beads were incubated for 2h at 4°C. Sonicated lysate with pre-incubated beads was added for an additional 4h. Beads were washed four times and GeneZol was added for RNA extraction. Serial dilutions of the 10% input cDNA (1:2, 1:10, 1:50, 1:250) were analyzed by real-time qPCR. The oligonucleotide sequences used are listed in SI Appendix, Table S4.

### Stable cell line establishment

pLenti CMV GFP Puro (Addgene, plasmid #17448) plasmid was cut with BsrgI and SalI restriction enzymes. Sequences of DNMT3B transcripts were isolated from pcDNA4-DNMT3B1 and pcDNA4-DNMT3B3ΔEx5 plasmids^33^ by PCR and assembled with pLenti construct using Gibson assembly (New England Biolabs) following the manufacturer protocol.

Lentiviruses were produced by transfection in the HEK293T packaging cell line using the PEI transfection reagent. Cells were transfected with each DNMT3B or GFP-only construct in addition to pCMV-dR8.2 dvpr (Addgene, plasmid #8455) and pCMV VSV-G (Addgene, plasmid #8454) lentiviral particles.

DNMT3B-null HCT116 cells^57^ were infected with viral particles and selected with Puromycin. Cells were incubated for 14 days to allow for stable expression and methylation dynamics.

### Bisulfite sequencing and analysis

Bisulfite conversion was performed using Zymo MethylGold kit following the manufacturer protocol. PCR reaction to isolate the D4Z4 region was done by using four previously described primer sets^58^ containing Illumina adaptor sequences at both the 5 ′ and 3 ′ ends. The PCR reaction was done with Taq Mix Red PCR MasterMix (Tamar). PCR products were cleaned and directly amplified using primers for the Illumina adaptor sequences followed by DNA sequencing. Methylation analysis was done using Biscuit (version 1.0.2). Reads were aligned using Biscuit align to the GRCh38 genome. The methylation level was estimated using Biscuit pileup with the -m 0 parameter, followed by the biscuit vcf2bed tool with the -t cg parameter. Otherwise, default parameters were used.

### Motif enrichment analysis

RNA-binding protein motif enrichment was performed using the SEA tool from the MEME suite (version 5.5.0)^59^. Enrichment analysis was performed for excluded exons sequences with included exons sequences as background, for human FSHD and mouse NSC results separately. FASTA files were generated using BEDTools getfasta tool (version 2.30.0)^60^ with hg38 and mm10 reference genomes, respectively. SEA analysis was performed using default parameters and a default E-value of ≤ 10, using the motif database: Ray2013 RNA (DNA-encoded)^61^.

## Acknowledgments

We thank Marnie Blewitt for valuable advice and Smchd1^MommeD1^ NSC RNA. We thank Abed Nasereddin and Idit Shiff from the Genomic Applications Laboratory, The Core Research Facility, The Faculty of Medicine - Ein Kerem, The Hebrew University of Jerusalem, Israel, for their professional advice and RNA-seq service. We thank Amos Tanay for DNMT3B-KO HCT116 cells. We thank Keith Robertson for pcDNA4-DNMT3B1, pcDNA4-DNMT3B3ΔEx5 plasmids. This work was supported by the Concern Foundation and AFHU / Boehm Foundation Young Professorship Award and the Alon Fellowship (YD).

