## Supplementary material for "SMCHD1 loss triggers DUX4 expression by disrupting splicing in FSHD2": Figures S1-S5

### Figure S1

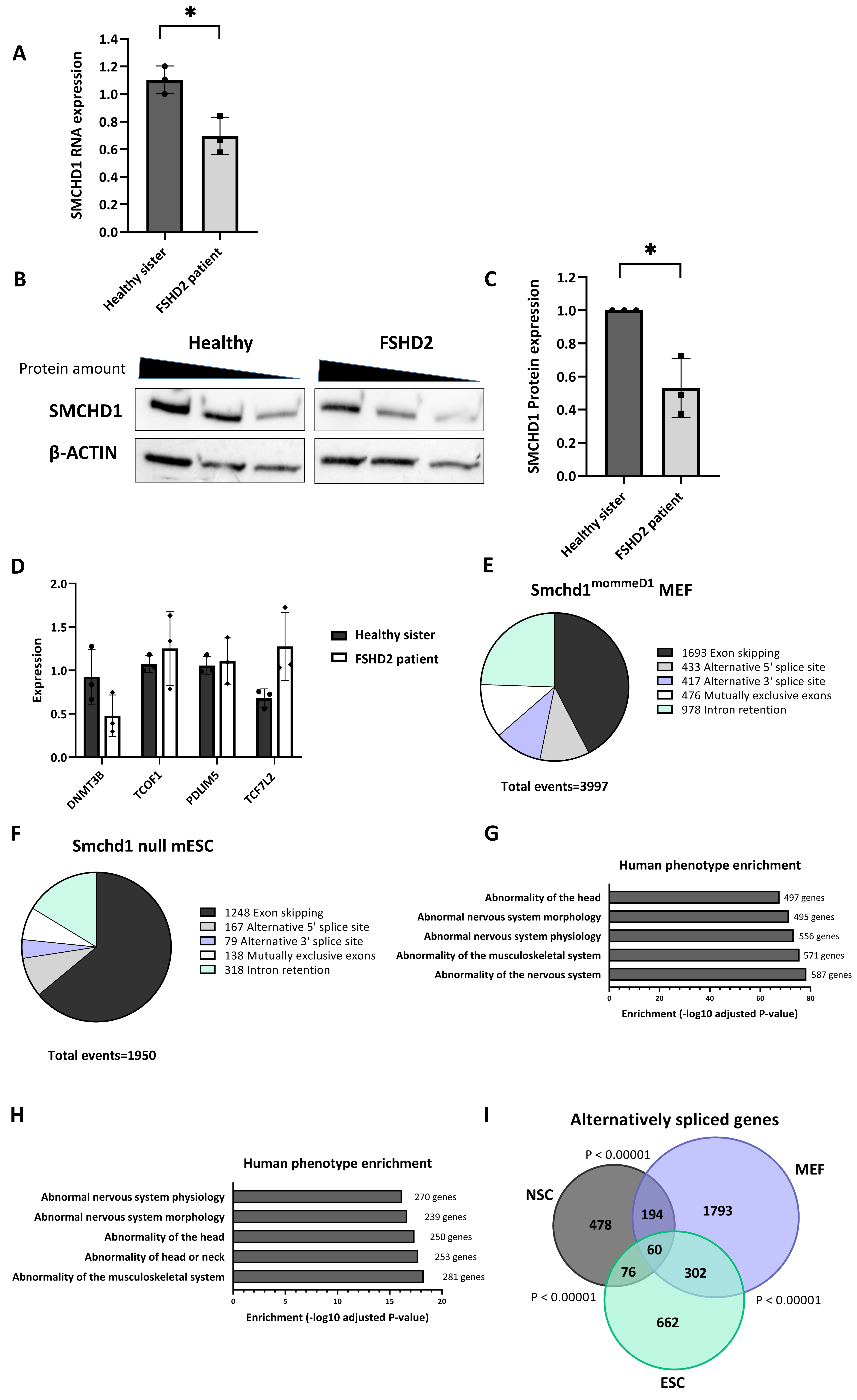


***Figure S1. A-D.*** RNA was extracted from lymphoblasts of an FSHD2 patient and her healthy sister and analyzed by real-time PCR for *SMCHD1* total mRNA amount relative to *CycloA* reference gene (**A**). Protein was extracted and Western blot was conducted with the indicated antibodies (**B**). Quantification of Western blot (C). RNA was extracted and real-time PCR was conducted to the indicated gene relative to *CycloA* reference gene (**D**). Values represent averages of three experiments ±SD, [* p<0.05]. ***E-H.*** RNA-seq of Smchd1$^{MommeD1}$ MEF **(E,G)** and Smchd1-KO mESC **(F,H)** samples was analyzed using rMATS (FDR<0.05). Proportion of each alternative splicing event is presented relative to WT , summary of significant alternative splicing events (FDR<0.05) from the five subtypes identified by rMATS in Smchd1$^{MommeD1}$ mice MEF RNA-seq data **(E-F)**. Top five significant events for human phenotype gene set, enrichment significance represented as –log10 adjusted P-value **(G-H)**. ***I.*** Venn diagram presenting the overlap between Smchd1 alternative splicing regulation in NSC, ESC and MEF of Smchd1 null or Smchd1-KO mice.

### Figure S2

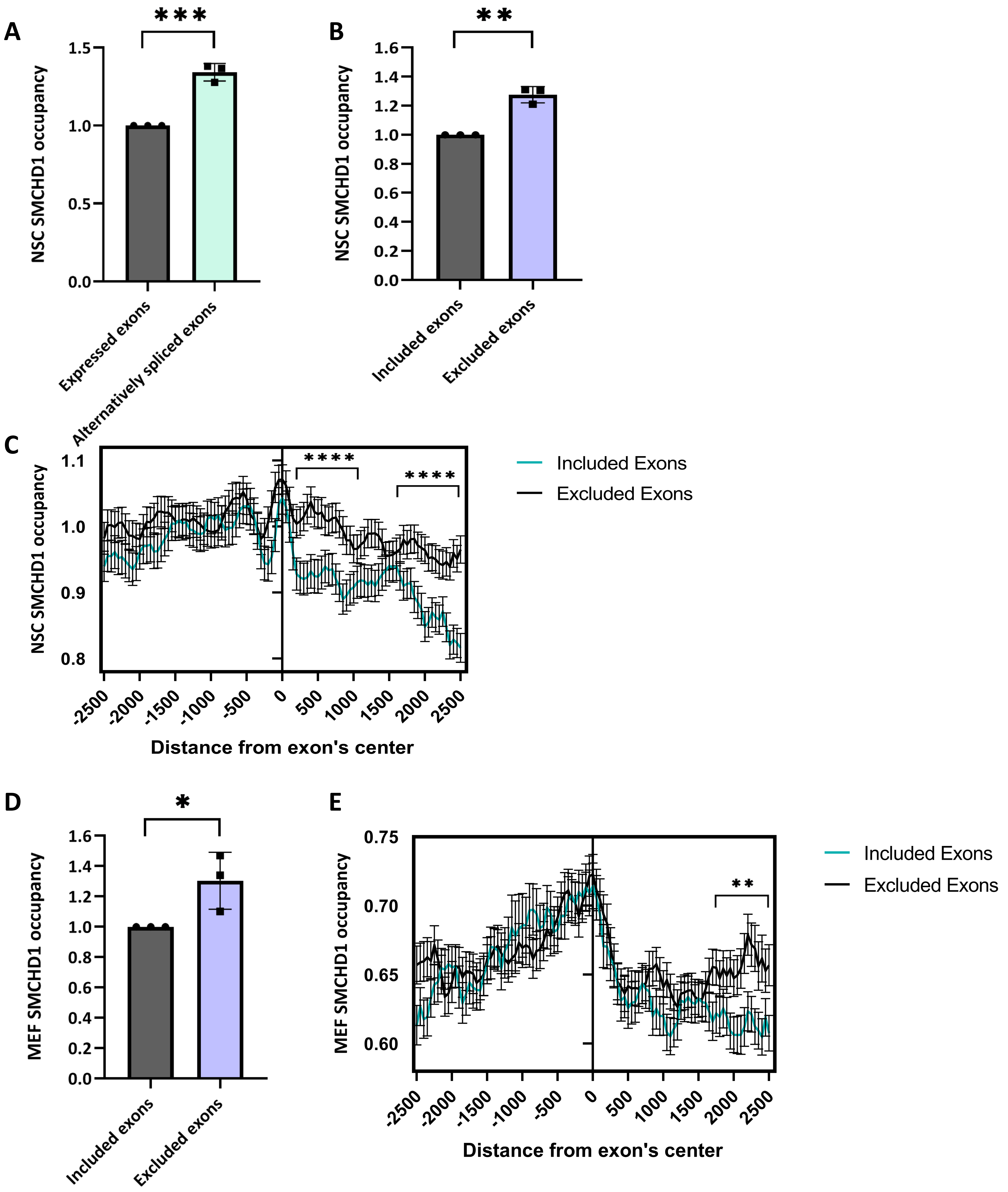


***Figure S2***

***A-C*** GFP ChIP-seq in primary NSCs with endogenous Smchd1-GFP fusion protein. Smchd1 occupancy analyzed as the number of Smchd1 peaks present within 5kb of alternatively spliced or expressed (TPM>1) exons **(A)** or within 5kb of differentially included or excluded exons in $Smchd1^{MommeD1}$ NSC **(B)** .Values represent averages of three ChIP-seq replicates from each cell type ±SD [* p<0.05; **p<0.01; ***p<0.001;]. Aggregation plot depicting the average normalized Smchd1 occupancy, at and near exons differentially included or excluded in $Smchd1^{MommeD1}$ mice. X axis represent bins of size 50 bp around the center of the exon [****p<0.0001] **(C)**. ***D-E***. SMCHD1 ChIP-seq in MEFs. SMCHD1 occupancy analyzed as the number of SMCHD1 peaks present within 5kb of included or excluded exons in $Smchd1^{MommeD1}$ MEF. Values represent averages of three ChIP-seq replicates from each cell type ±SD [* p<0.05; **p<0.01; ***p<0.001;] (A). Aggregation plot depicting the average normalized Smchd1 occupancy, at and near exons differentially included or excluded in $Smchd1^{MommeD1}$ MEF. X axis represent bins of size 50 bp around the center of the exon [**p<0.01].

### Figure S3

A

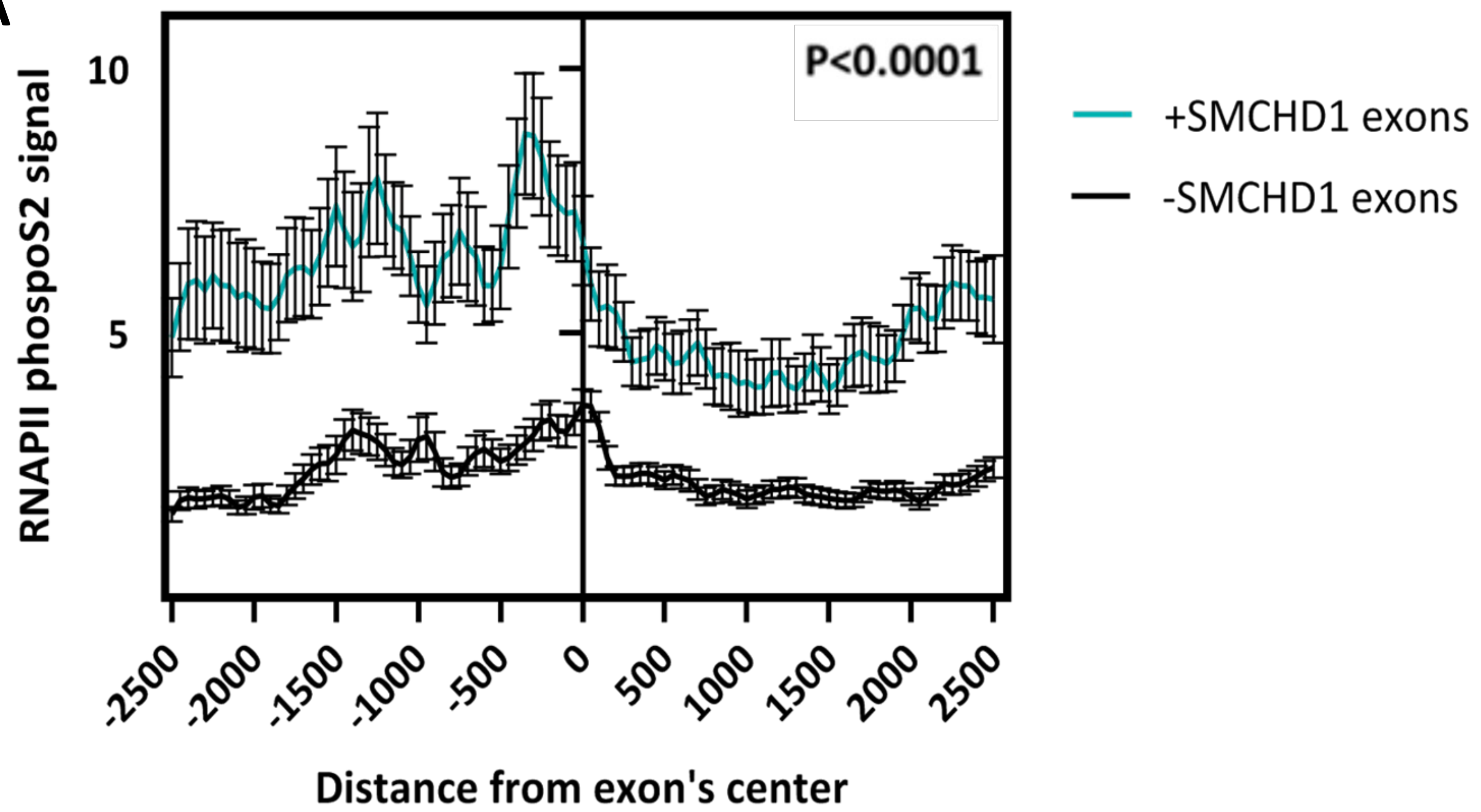


B

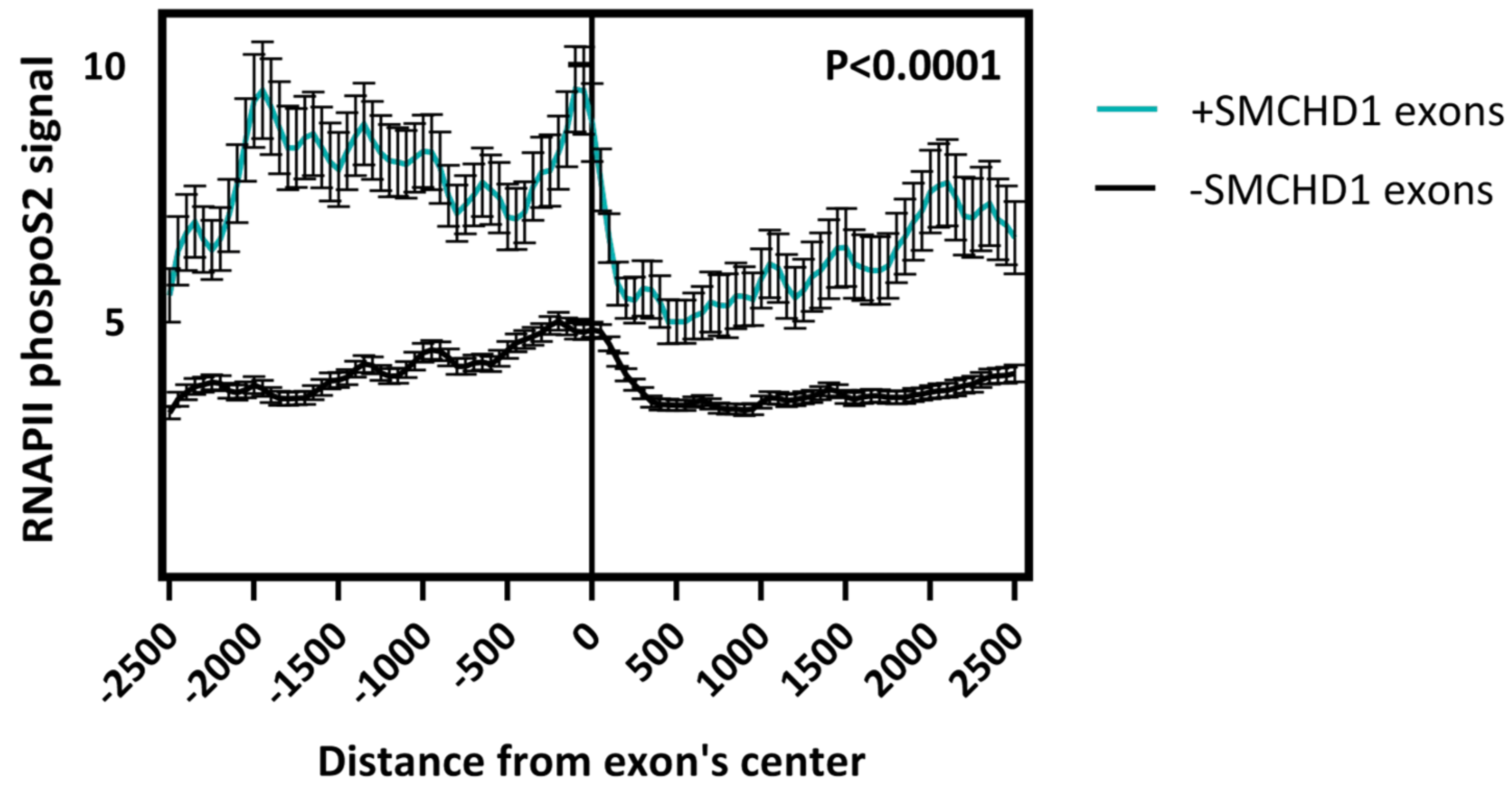


***Figure S3***

**A-B.** Aggregation plot depicting the average normalized phospho-Ser2 levels of RNAPII at and near alternatively spliced exons differentially bound by Smchd1 in NSC (A) or MEF (B). X axis represent bins of size 50 bp around the center of the exon .

### Figure S4

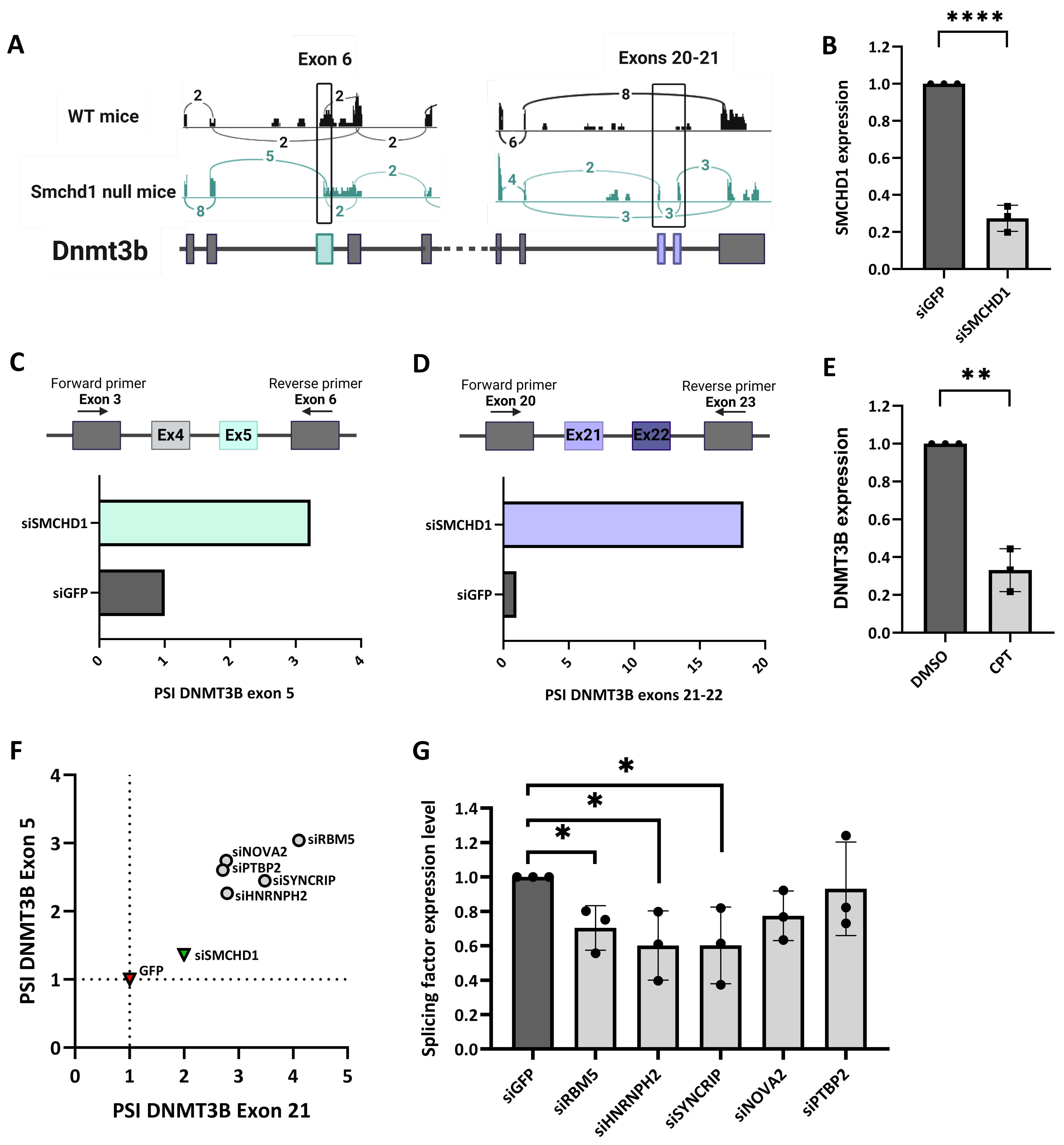


***Figure S4***

***(A)*** RNA was extracted from three Smchd1$^{MommeD1}$ mice NSC samples and two WT mice NSC samples. RNA-seq was conducted and analyzed using rMATS (FDR<0.05). Genome browser view of the Dnmt3b alternatively spliced junctions presented by sashimi plots, arcs denote splice junctions quantified in spanning reads. ***(B-D)*** HCT116 cells were transfected with siRNA targeting SMCHD1 or GFP as negative control. Total RNA was extracted and analyzed by real-time PCR for SMCHD1 total mRNA amount relative to *CycloA* reference gene. Values represent averages of three experiments done in triplicates ±SD normalized to negative control (siGFP) [****p<0.0001] (paired Student's t-test) **(B)**. Semi quantitative PCR was conducted for exons 4-5 **(C)** and exons 20-21 **(D)** using custom primers as described. PSI is calculated as the included product amount relative to the excluded product, and normalized to siGFP as a negative control. ***(E)*** HCT116 cells were treated with 6uM of CPT or DMSO as negative control for 6 hr. Total RNA was extracted and analyzed by real-time PCR for total DNMT3B mRNA amount relative to *CycloA* reference gene. Values represent averages of three experiments done in triplicates ±SD normalized to negative control (DMSO) [**p<0.01] (paired Student's t-test). ***(F)*** HCT116 cells were transfected with siRNA targeting 71 human splicing factor, SMCHD1 as a positive control and GFP as negative control. Total RNA was extracted and analyzed by real-time PCR for DNMT3B exon 5 and exon 21 relative to DNMT3B total mRNA amount. PSI was calculated as DNMT3B exon inclusion/DNMT3B total mRNA and normalized to negative control (siGFP). Values represent averages of two experiments ±SD. Negative control (siGFP) PSI is represented by the dotted line at 1. ***(G)*** HCT116 cells were transfected with siRNA targeting each splicing factor hit indicated or GFP as a negative control. Total RNA was extracted and analyzed by real-time PCR for splicing factor total mRNA amount relative to *CycloA* reference gene. Values represent averages of three experiments done in triplicates ±SD normalized to negative control (siGFP) [*p<0.05] (paired Student's t-test).

### Figure S5

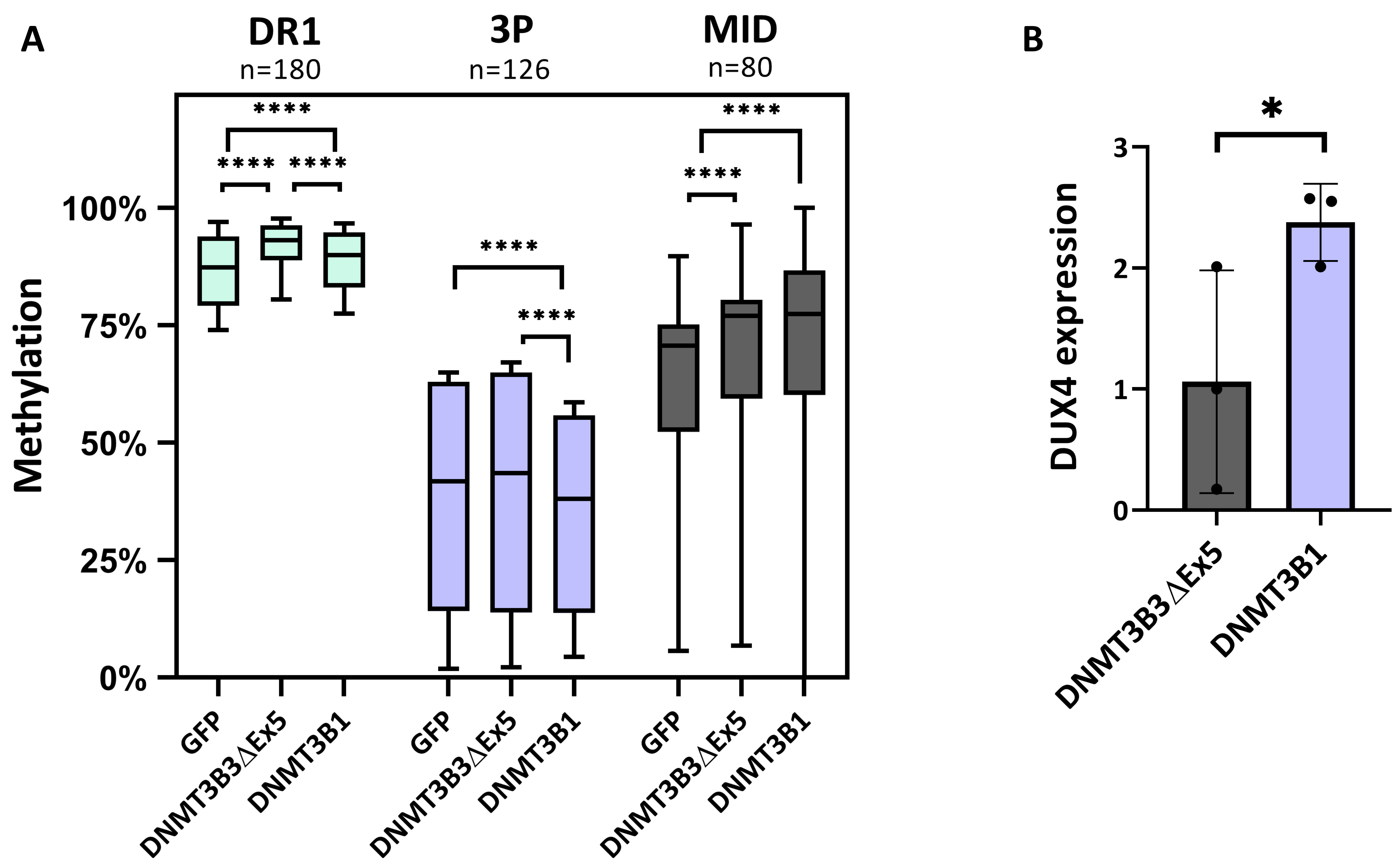


***Figure S5***

***(A)*** DNMT3B null HCT116 cells were infected with empty-GFP, GFP-DNMT3B3ΔEx5 or GFP-DNMT3B1 lentiviruses. DNA was isolated and bisulfite converted. PCR was conducted in three locations along the D4Z4 region (see methods) and products were sequenced. Methylation level in each CG was assessed using Biscuit. Methylation level is presented at three different regions of D4Z4 region (see methods). ***(B)*** RNA was extracted from cells with the DNMT3B3ΔEx5 and DNMT3B1 isoforms and real-time PCR was conducted. DUX4 mRNA level was quantified relative to *CycloA* reference gene. Values represent averages of three technical replicates ±SD; [* $p<0.05$] (Student's t-test).
